# Statistical inference of protein structural alignments using information and compression

**DOI:** 10.1101/056598

**Authors:** James H. Collier, Lloyd Allison, Arthur M. Lesk, Peter J. Stuckey, Maria Garcia de la Banda, Arun S. Konagurthu

## Abstract

Structural molecular biology depends crucially on computational techniques that compare protein three-dimensional structures and generate structural alignments (the assignment of one-to-one correspondences between subsets of amino acids based on atomic coordinates.) Despite its importance, the structural alignment problem has not been formulated, much less solved, in a consistent and reliable way. To overcome these difficulties, we present here a framework for precise inference of structural alignments, built on the Bayesian and information-theoretic principle of Minimum Message Length (MML). The quality of any alignment is measured by its explanatory power - the amount of lossless compression achieved to explain the protein coordinates using that alignment. We have implemented this approach in the program `MMLigner` http://lcb.infotech.monash.edu.au/mmligner to distinguish statistically significant alignments, not available elsewhere. We also demonstrate the reliability of `MMLigner`’s alignment results compared with the state of the art. Importantly, `MMLigner` can also discover different structural alignments of comparable quality, a challenging problem for oligomers and protein complexes.

---

An alignment of two (or more) proteins is an assignment of a one-to-one correspondence between subsets of their amino acid residues. An alignment can be inferred from amino acid sequence information (yielding a sequence alignment) or using the information of three-dimensional atomic coordinates of residues (yielding a structural alignment). As proteins diverge during evolution, structure changes more conservatively than sequence.^1–3^. Therefore, it becomes necessary to appeal to structure when inferring a trustworthy relationship between distantly-related proteins^4,5^.

Protein alignment is essential for many areas of research in molecular biology. The purpose of an alignment is to identify similarities between two or more proteins, both in terms of sequence and in terms of structure. Relevant problems include identifying homologous amino acids, *i.e.*, residues encoded by nucleotides at equivalent positions in genome sequences,^5 6^ understanding the evolution of protein domains, homology modelling of proteins of known amino-acid sequence but unknown structure, and supporting experimental protein crystal structure determination using molecular replacement techniques^4^.

The structural alignment problem is the identification of a subset of residues with matching three-dimensional structural contexts. Most homologous proteins preserve a *common core* within their three-dimensional structure.^1^ This core comprises one or more regions of residues that retain the same topology of their folding patterns, varying only in their spatial and geometric details.^5^ Peripheral regions outside the common core tend to refold entirely, and cannot be meaningfully aligned. Although pairwise sequence alignment cannot identify such regions, structural alignment can - and must.

The structural alignment problem can be cast as an optimisation problem. This requires: (1) a *measure of quality* to assess any proposed alignment between proteins, and (2) a *search method* to find an optimal alignment under the stated measure. Over the past five decades, very many structural alignment methods have been proposed, differing mainly in how they assess structural alignment quality.^7^ By one estimate, the number of new structural alignment methods is doubling roughly every five years^8^.

Recent comparative studies have exposed disagreements among the results produced by different currently available structural alignment methods.^7–10^ This disagreement is traceable to the lack of a systematic framework for formulating the structural alignment problem, especially in the ambiguity in defining a rigorous measure of alignment quality to be optimised. (This deficiency stands in stark contrast with the progress achieved in the closely related field of protein sequence alignment, where research has produced rigorous statistical framework for sequence comparison.^11–13^ Largely due to the standardisation of the framework, the algorithmic development for computing sequence alignments has ‘reached a point of diminishing returns’^14^).

Providing a rigorous framework for evaluating structural alignments is likely to rectify the lack of consensus that has beset the field of structural alignments. To achieve this goal, we present a novel statistical measure to assess structural alignment quality.^15^ This measure does not suffer from the ambiguities that have been so problematical in other approaches.

Our measure of alignment quality is built on the rigorous framework of Minimum Message Length (MML) criterion.^16,17^ MML provides a general purpose Bayesian framework for statistical inductive inference, useful to discriminate between competing hypotheses on observed data.^17,18^ In this context, any structural alignment is treated as a hypothesis of the structural relationship between two proteins, expressed as a one-to-one, order-preserving, correspondence between subsets of residues. Specifically, each alignment hypothesis constitutes an attempt to explain the observed C*_α_* coordinates. The explanatory power of each alignment is quantified, using principles of information theory, as the amount of lossless compression gained in encoding the C*_α_* coordinate data of the aligned proteins.

This measure can be thought of as a communication between an imaginary transmitter-receiver pair. The quality of any structural alignment is measured as the message length (in bits) required to transmit the C*_α_* coordinates using the alignment hypothesis, allowing it to be received and decoded losslessly. This measure is backed by mathematical rigour and exhibits important statistical properties, including a natural null hypothesis test for assessing the statistical significance of any proposed alignment (**Methods** and **Supplementary Notes S1-S3**).

We describe our algorithm `MMLigner`, designed to search for, and report, statistically significant structural alignments. This algorithm uses, as its measure of alignment quality, the message length (in bits) required to transmit protein C*_α_* coordinates using the alignment hypothesis. The code we developed is available publicly under an open source (GPL) licence at http://lcb.infotech.monash.edu.au/mmligner and as **Supplementary Software**.

Comprehensive benchmarking demonstrates the effectiveness of our new approach to protein structural alignment and compares it with that of the available state of the art approaches. Importantly, MMLinger can also identify and evaluate different alignments for the same pair of protein structures. This is especially useful when aligning oligomers and protein complexes, where conventional structural alignment techniques are generally inadequate^10,19^.

## Methods

### Problems with defining an optimal alignment

The difficulty with many structural alignment methods arises from the problem of specifying precisely what needs to be optimized. There is a conflict - requiring some kind of compromise - between two key alignment criteria that besets the current alignment programs: *coverage* and *fidelity*. Conventional methods typically approximate coverage as the size of the alignment: the number of correspondences plus, in some cases, the number of residues in the unaligned regions. Fidelity measures the spatial or geometric similarity of aligned residues, commonly approximated either by the root-mean-square deviation (RMSD) computed after the best rigid-body superposition, or by pairwise Euclidean distance profiles.

Different compromises between the weighting of maximum coverage and minimum RMSD in conventional structural alignment approaches are a major source of the inconsistencies in their results. For example, suppose we are aligning two 300-residue proteins. It may be possible to find one alignment of 200-residue substructures of the proteins, with RMSD 2.5 Angstroms (Å) with one weighting, and another alignment of 100-residue substructures with RMSD 2.0 Å with a different weighting. It is this difficulty that led Chothia and Lesk to define the core of a pair of structures to choose the subsets of structures to align^1^.

### Principles and main features of the Minimum Message Length framework

Inductive inference aims at identifying *hypotheses* that best explain observed data.^17^ In general, a complex hypothesis can explain (or fit) a greater variety of observed data than a simpler hypothesis.^15^ Therefore, choosing the best hypothesis for any inference problem involves a necessary tradeoff between hypothesis complexity and its fit with the observed data. Structural alignment can thus be seen as an instance of the general class of inference problems. As suggested earlier, an alignment is a hypothesis that attempts to *explain* the relationship between two proteins, given as the observed data the coordinates of the C*_α_* atoms. In conventional statements of the structural alignment problem, the aforementioned complexity versus fit trade-off is handled using surrogate, easily-computable criteria. (Traditionally, coverage is used as a proxy for hypothesis complexity, while RMSD is used as a proxy for fit with observed data).

The field of statistical learning and inference provides rigorous approaches to address this problem systematically. In the early nineteen sixties several landmark papers proposed links between inductive inference and information theory.^16,20,21^ The MML principle provided the first practical information-theoretic criterion for hypothesis selection based on observed data. MML is used here to assess structural alignment quality and to differentiate reliably among competing alignments. *This solves the problem of coverage-versus-RMSD tradeoff by allowing both to affect the Message Length, minimization*.

How is information content measured? Claude E. Shannon in his seminal work on the mathematical theory of communication provided the means to quantify information content: It is the length (in bits) of the shortest message needed to communicate an event losslessly between an imaginary *transmitter-receiver* pair. Just as it requires one bit to report the result of an (unbiased) coin toss, if some event *E* has the probability Pr(*E*), the Shannon Information content (represented as *I*(*E*)) of that event is denoted as — log_2_(Pr(*E*)) (measured in bits)^22^.

Consider a *hypothetical* communication process between the *transmitter-receiver* pair. The transmitter has the protein coordinate data of two structures, *S* and *T*, and wants to communicate both sets of C*_α_* coordinates to the receiver. One approach is to transmit *S* and *T* independently, by assuming they are *unrelated* to each other. We call this form of encoding the *null model* message. The length of the null model message for transmitting both *S* and *T* is denoted as:

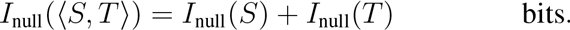

An alternative approach to transmit *S* and *T* is to exploit the structural *relationship* (if any) between them using an alignment, denoted as 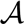. We call this form of encoding the alignment the *relationship model* message. Intuitively, the relationship model message can be decomposed into two parts. In the first part the transmitter communicates the information of the alignment 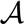 as a string over `match (m), insert (i)`, and `delete (d) states`. Receiving this information, the receiver can decode and reconstruct the alignment between protein structures *S* and *T*, but has no knowledge (yet) of their coordinates. Therefore, the second part encodes the coordinate information of *S* and *T* using the alignment 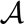.

The total message length of the relationship model message (or `I-value`) for transmitting both *S* and *T* given 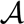 can be represented as

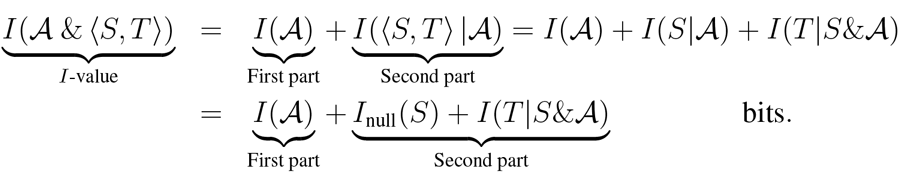

The above equation is the result of Bayes’ theorem restated in Shannon’s information-theoretic terms (**Supplementary Note S1**). We shall denote 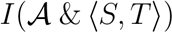 by `I-value` (**Supplementary Notes S2-S3**).

If the proposed alignment relationship, 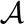, is a poor one (few residues aligned, large structural deviations), then the encoded relationship model message will be inefficient, making `I-value` long. Alternatively, if the alignment relationship is a good one (many residues aligned, small structural deviations), then the relationship model is efficient, yielding a more concise message; that is, a smaller `I-value`. Thus, `I-value` forms an excellent measure to assess structural alignment quality. An optimal alignment is one with the shortest total message length; that is, with the lowest value (in bits) for `I-value`.

Furthermore, the difference between the null and the relationship model message lengths provides a measure of the statistical significance of the alignment hypothesis (**Supplementary Note S1**):

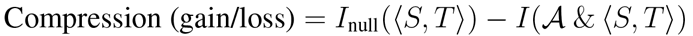

If the above difference is positive, that is, if relationship model message gains compression over the null model message, then the alignment is considered significant. If, on the other hand, the relationship model message is more verbose (*i.e*., has a greater length) than the null model message, then that alignment is rejected. This ability to detect statistically significant alignments is unique to our method.

### MMLigner algorithm

Having defined `I-value` as the quantity to be optimised, we must supply a search strategy to achieve this. The search is carried out in two phases. (**Supplementary Note S4**.)

In the first phase, crude-but-effective structural alignments are rapidly generated, to act as seeds for refinement using the `I-value` measure. To this end; we first identify continuous fragments from the two structures that superpose well (below a threshold value of RMSD). We call this the library of maximal fragment pairs (MFPs). Each MFP contains one region from each protein. This library of MFPs is filtered efficiently^23^ by exhaustively superposing (non-overlapping) pairs of MFPs (then triples), and retaining only those MFPs for which there are partners such that the joint superposition of the MFP and its partners have an RMSD lower than a fixed threshold. This results in a drastically-pruned library of MFPs, in which each MFP can be jointly superposed with at least two other (non-overlapping) MFPs in the set. Next, these filtered MFPs are efficiently clustered. For each cluster, a dynamic programming algorithm^5^ is used to generate a crude seed structural alignment.

In the second phase, the seed alignment for each cluster is iteratively refined over an Expectation-Maximization approach using the `I-value` criterion (**Supplementary Note S3**). The refinement is carried out via an ensemble of alignment perturbations while keeping track of the amount of compression achieved compared to the null model (**Supplementary Note S2**).

This search is illustrated in Fig. 1 using an example pair of structures: the chain A of pig heart, GTP-specific succinyl-CoA synthetase (`1EUD-A`, 306 residues) and the chain A of glutamate mutase from *Clostridium cochlearium* (`1CCW-A`, 137 residues). Note that this part has two distinct structural alignments of comparable quality (Table 2).

## Results

### Software Benchmarking

We benchmark the performance of `MMLinger` and compare it with five widely used alignment programs (`DALI`^24^, `LGA`^25^, `FATCAT`^26^, `CE`^27^ and `TM-Align`^28^). A dataset of protein structural domains is taken from the SCOP database.^29^ This results in a random selection of 2500 domain pairs, with variable closeness of structural relationships, according to the SCOP hierarchy: Family, Superfamily, Fold, Class and Decoy (in equal proportions) (**Supplementary Note S5**). By a decoy we mean a protein that is not in the same Class as another, to which we attempt an alignment as a control. `MMLinger` and other alignment programs are used to align these structural pairs, each generating 2500 alignments (i.e., 500 alignments per each SCOP group).

**Figure 1:**
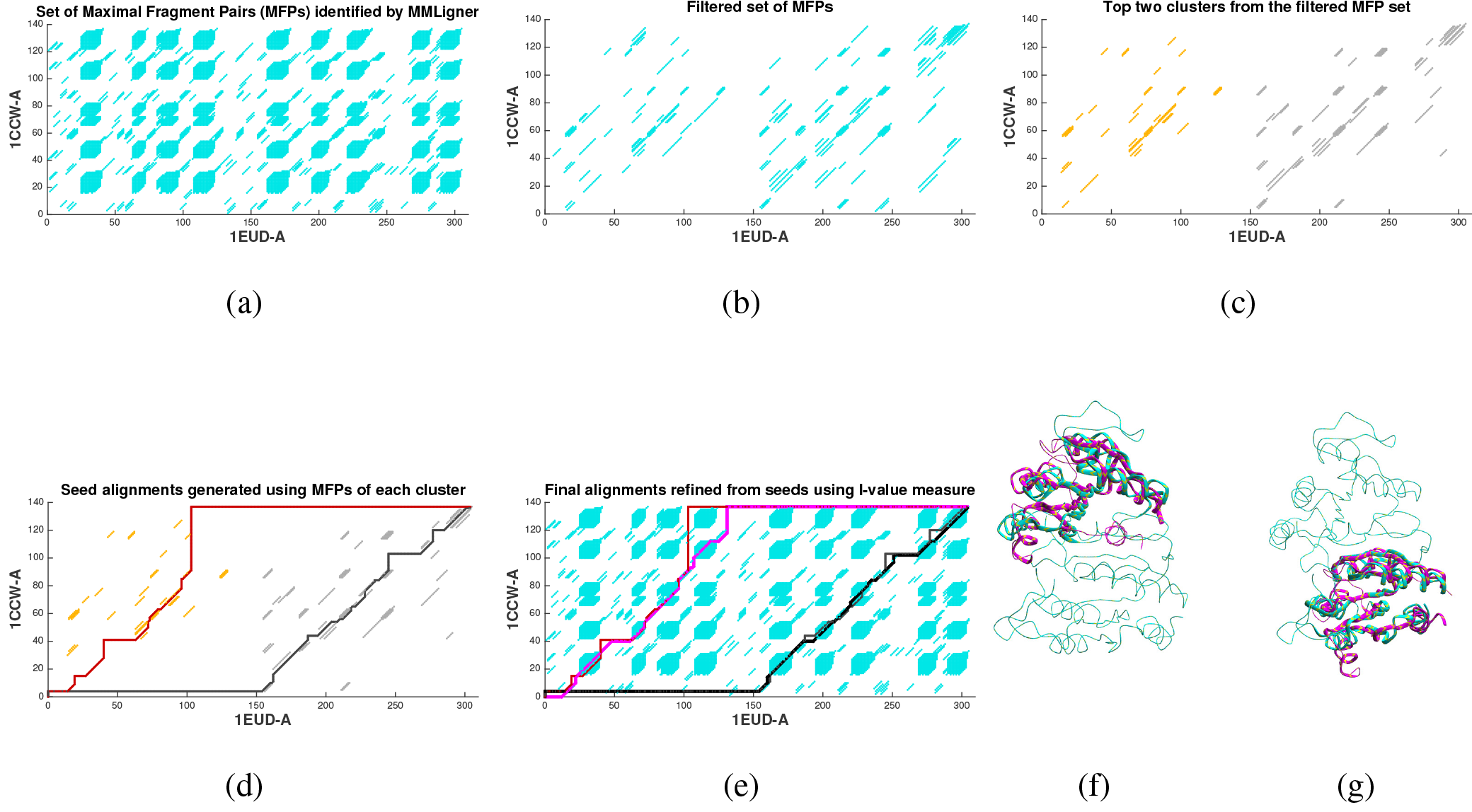
Illustration of `MMLinger`’s search using the example of `1EUD-A` and `1CCW-A`. (a) The full library of maximal (well-superposable) fragment pairs (MFPs) identified by `MMLinger` shown as a 2D dot plot. That is, each blue dot within a blue region corresponds to a residue `1EUD-A` (on the x-axis) and a residue in `1CCW-A` (on the y axis). A blue point at this intersection indicates that there is an MFP in both proteins that includes the pair of residues corresponding to the projections of the points on the axes. (b) Filtered set of MFPs, where each listed MFP jointly superposes with at least two other (non-overlapping) MFPs in the set. (c) Clustering MFPs into groups that fit consistently. Top two clusters are shown here. (d) Determination of seed alignments using the MFPs in each cluster. The seed alignment is shown as a path on the 2D dot plot. (e) Final `MMLinger` alignments after iterative refinement of the seed alignments using `I-value` measure. The seed alignment is also juxtaposed for reference. (f-g) Superpositions of `1CCW` on `1EUD`, based on the two `MMLinger` alignments. (Table 2; **Supplementary Movie S1(a)-S1(b))**.

We report in Figure 2 both conventional measures of alignment quality - coverage and RMSD - and also information-based complexity 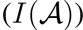 and compression (in terms of `I-value`) relative to the null message. Focusing first on the alignment coverage measured as the number of residue-residue correspondences (Fig. 2, first column), we expect the coverage to decrease as the structural similarity decreases. This is indeed the case for all programs tested, with `MMLinger` showing the most pronounced decrease, particularly for Family, Superfamily and Fold. (For a large majority of structural pairs in the Class and Decoy sets, `DALI` does not return any alignment. We interpret these cases as the structures being entirely unaligned, yielding zero coverage and RMSD.).

The median coverage that `MMLinger` produces for alignments of pairs of proteins in the same Class group is approximately 25 correspondences, and the value for alignments with Decoys is zero. Inspecting these groups manually when alignment coverage is greater than zero, we find that `MMLinger` typically aligns up to two distinct supersecondary structures for the Class dataset, and up to one supersecondary structure for the Decoy set. Comparing the plots of different alignment programs, the median coverage values for `DALI`, `TM-Align` and `LGA` are similar to each other, and are higher than those reported by other programs: Although `MMLinger`’s median coverage is only slightly smaller, those of `CE` and `FATCAT` are significantly smaller, indicating they are the most conservative of all the programs. Conservative, in this context, means biased towards smaller alignments (lower coverage). But coverage alone is not meaningful without also considering the structural deviation of the aligned residues.

**Figure 2:**
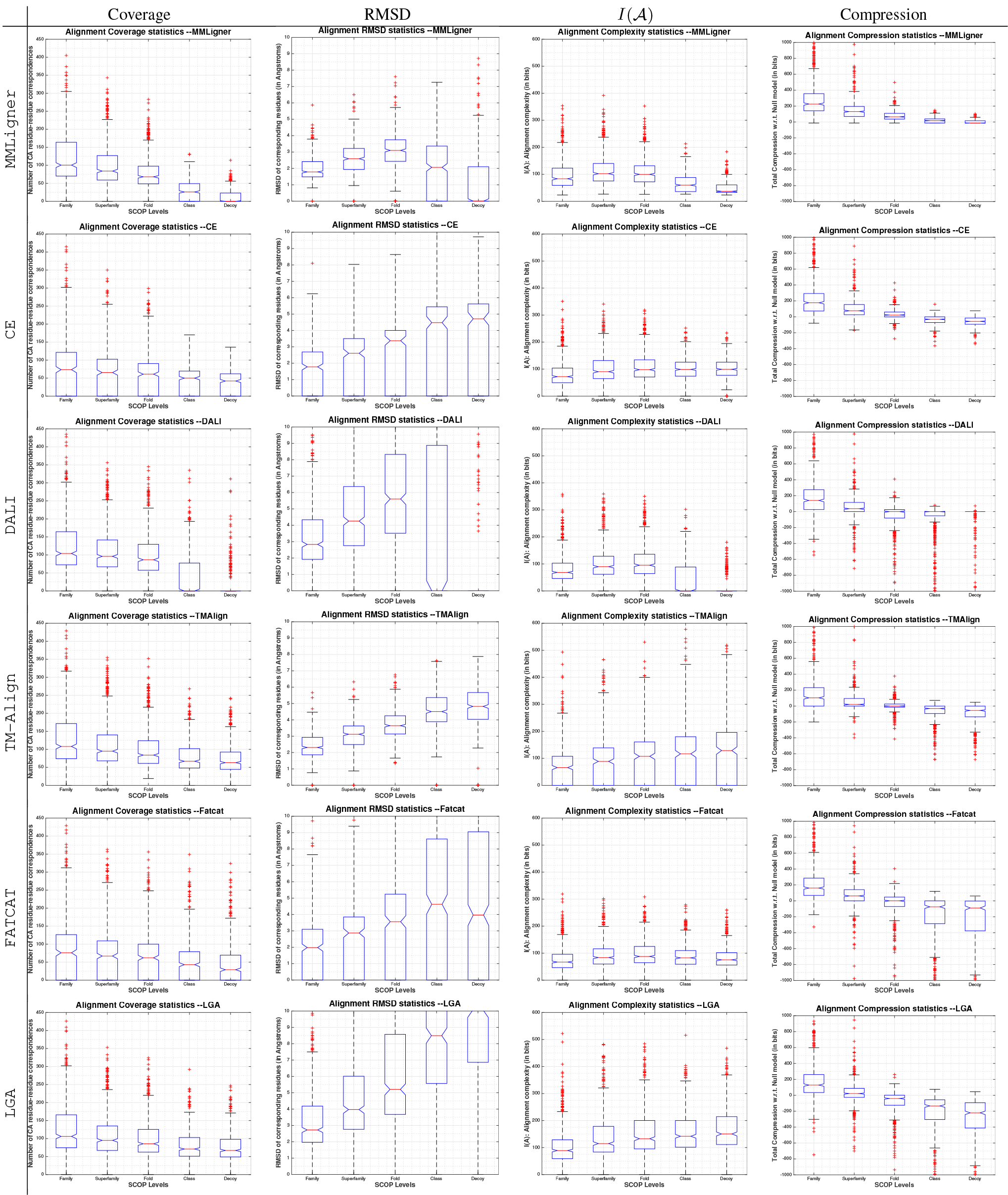
Notched Box-Whisker plots displaying the distribution of values from various criteria across 500 x 5 = 2500 alignments generated by various alignment programs. Rows indicate the alignment program used: `MMLinger`, `CE`, `DALI`, `TM-Align`, `FATCAT`, and `LGA` (in this order). Columns indicate various alignment criteria: Coverage (number of one-to-one correspondences), RMSD (after best rigid-body superposition), Alignment complexity 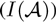 and Compression (with respect to the null model message length). Note that the ordinate scale for each of the four criteria has been constrained to be equal throughout any column, so that the plots across various alignment programs are visually comparable.

Focusing on alignment fidelity measured by the RMSD of the best superposition (Fig. 2 second column), we expect the RMSD first to increase from Family to Superfamily and Fold, as the structural relationship progressively diverges, and then to decrease as the coverage falls due to the proteins’ no longer having any major structural relationship. This is the case for `MMLinger`, `DALI` and, to some extent, `FATCAT` (Fig. 2 second column, first/third/fifth rows). However, the other aligners retain an increasing trajectory, as their scoring functions do not inherently compromise between the conflicting objectives arising from simultaneously maximising coverage and minimising RMSD.

Comparing the graphs of the results of the different alignment programs, `MMLinger`’s median values for RMSD is the lowest across the Family, Superfamily and Fold groups, followed by `CE` and `FATCAT`; with `DALI` and `LGA` yielding alignments with poor fits. The relatively large coverage values imply that `DALI` and `LGA` have a tendency to over-align residues at the cost of their fit. This behaviour has been observed upon careful manual study of many alignments and superpositions. `TM-Align`’s RMSD profile, although poorer than those of `MMLinger`, `CE`, and `FATCAT`, remains reasonably well behaved for the Family, Superfamily and Fold groups. However, upon manual inspection, `TM-Align` tends to align greedily single and pairs of residues when they randomly appear spatially proximal, yielding more complex alignments (Fig. 2 third column).

Analysing the spread of the interquartile (IQR) regions across various alignment programs for both coverage and RMSD, we find that `MMLinger` is the best behaved overall across these two measures (Fig. 2, first/second columns). The compactness of these boxes indicates `MMLinger`’s consistency over other programs to identify meaningful structural alignments. The non-overlap of the notches (that signifies the confidence interval) in the box plots of `MMLinger` suggest that the medians are statistically different. No other program produces non-overlapping notches between boxes across all groups for the coverage statistics.

Comparing the complexity 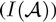 of various alignments (Fig. 2, third column), we notice that the median lines for `MMLinger` across the five groups show a distinct concave trajectory. As the structures diverge from Family to Superfamily and Fold, the complexity of the relationship increases, and so does the alignment explanation length, the first part of the message. Then 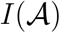 starts to decrease for the alignments in the Class and Decoy groups since their relationship (if any) remains simple, and hence can be stated concisely. Only `DALI` among the other programs shows this trajectory. However, its concavity is an artifact of the large majority of the alignment from the Class and Decoy groups going unreported (treated here as zeros). For other programs, 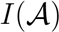 median lines show either an increasing trend, or stay relatively flat.

The total compression gained in bits by the various alignments (Fig. 2, fourth column), shows that `MMLinger` gives the best compression overall. This is not surprising, as `MMLinger` attempts to optimise this criterion. Nevertheless, `I-value` has a utility beyond comparing competing alignments of the same pair of structures: it identifies which alignments are significant. In this respect, `I-value` goes beyond conventional measures that build on coverage and fidelity. The horizontal line at 0 bits is the discriminating line for statistical significance in the information-theoretic framework. When compression is positive (above the zero line), the alignment provides a more concise lossless explanation of the coordinates of the structural pair than their null model explanation. Such an alignment should be accepted as significant (**Supplementary Note S1**). On the other hand, if the compression is negative (below the zero line) the lossless explanation using the alignment relationship is longer than the null model message length. Such an alignment should be rejected.

For `MMLinger`, the boxes (hence their corresponding alignments) for Family, Superfamily and Fold are always above the zero line, suggesting that these alignments are significant, as measured by our lossless compression based framework (Fig. 2, first row, fourth column). Surprisingly, except for `CE`, other programs produce, on average, alignments for the SCOP Fold group which are below the zero line, and which, although they may be optimal with respect to some operative measure of alignment quality, are not statistically significant. `MMLinger`’s first quartile line for the Class group and the third quartile line for the Decoy group sits close to zero. For the pairs in the Class and the Decoy groups that yield positive compression, many alignments involved alpha-hairpins covering over 20 or more residues. (MMLigner provides an additional option, handled as a postprocess, to reject any alignment with low coverage that aligns supersecondary structures. The results/box-plots of such runs are shown in **Supplementary Note S6**). As mentioned before, `DALI` does not return any alignment for most of the Class and almost all of the Decoy sets, explaining their quartile lines.

In summary, the performance of `MMLinger` is consistently reliable, compared to all other programs considered. True to the MML framework, which relies on achieving an accurate tradeoff between hypothesis (here, alignment) complexity and fidelity to the observed data (here, the lossless explanation of structural coordinates), `MMLinger` consistently identifies statistically significant alignments when they exist (and not when they don’t), avoids pairing up spurious correspondences, and favours simple alignments over complex ones.

### Identification of alternative structural alignments

MMLigner’s ability to identify alternative alignments for the same given structural pair is demonstrated in the following two case studies.

*Aligning 2SAS-A with 1JFJ-A:* Consider the *α* chains (chain A) of the pair of calcium-binding proteins from *Branchiostoma lanceolatum* (wwPDB entry: `2SAS` chain A with 134 residues) and *Entamoeba histolytica* (wwPDB entry: `1JFJ` chain A with 185 residues). SCOP^29^ classifies the corresponding domains, `d2sasa`_ and `d1jfja`_, within the Calmodulin-like family, suggesting a close evolutionary relationship. Each of these domains contains four EF-hand (helix-loop-helix) motifs. Two EF-hands pair up near the N-terminus and the other two pair up near the C-terminus in each domain. However, these EF-hand pairs have markedly different topologies: they are flexed compactly in `d2sasa`_, while they remain relaxed in `d1jfja`_. `MMLinger` produces four distinct alignments that rate as statistically significant, based on the compression those alignments achieve over the null model message length. These four alignments correspond to the four possible ways in which the EF-hand pairs can match with each other. Fig. 3 shows the alignments found by `MMLinger` and their corresponding least-squares superpositions.

Most other alignment programs return just one structural alignment of the four possible alignments. This is shown in Table 1, which gives the traditional alignment statistics, coverage and RMSD, generated by `MMLinger` and other popular programs for comparison. Also, shown in Table 1 are the information measures of alignment complexity and compression with respect to the null model message length (both in bits). `DALI`^24^ is the only other program besides `MMLinger` that reports all four alignments. However, only two of those have reasonable RMSD values. The other programs, `LGA`^25^, `FATCAT`^26^, `CE`^27^ and `TM-Align`^28^, produce only one of the four alignments, with varying alignment quality based on their Coverage and RMSD values.

*Aligning `1EUD-A` with `1CCW-A`:* Consider now the alternative alignments for the pair 1EUD (chain A) and 1CCW (chain A) discussed earlier when describing `MMLinger`’s algorithm (Fig. 1(e)-(f)). These correspond to the *α* chains (chain A) of proteins pig Succinyl-CoA synthetase (wwPDB entry: `1EUD-A` with 306 residues) and Glutamate mutase from *Clostridium cochlearium* (wwPDB entry: `1CCW-A` with 137 residues). SCOP dissects the *α* chain of Succinyl-CoA synthetase into two domains, d1euda1 (130 residues) and `d1euda2` (176 residues), and classifies them under different folds. The N-terminal (CoA-binding) domain, d1euda1, is classified as an NAD(P)- binding fold, while the C-terminal domain, `d1euda2`, is classified as a Flavodoxin-like fold. On the otherhand, SCOP assigns the *α* chain of Glutamase mutase to a single domain, `d1ccwa_`, classified under Flavodoxin-like fold, the same as for `d1euda2`. Interestingly, the two domains of `1EUD-A` share self-similarity and, hence, it is possible to align the Glutamase mutase domain to either the N-or C-terminal domains of Succinyl-CoA synthetase.

**Table 1:**
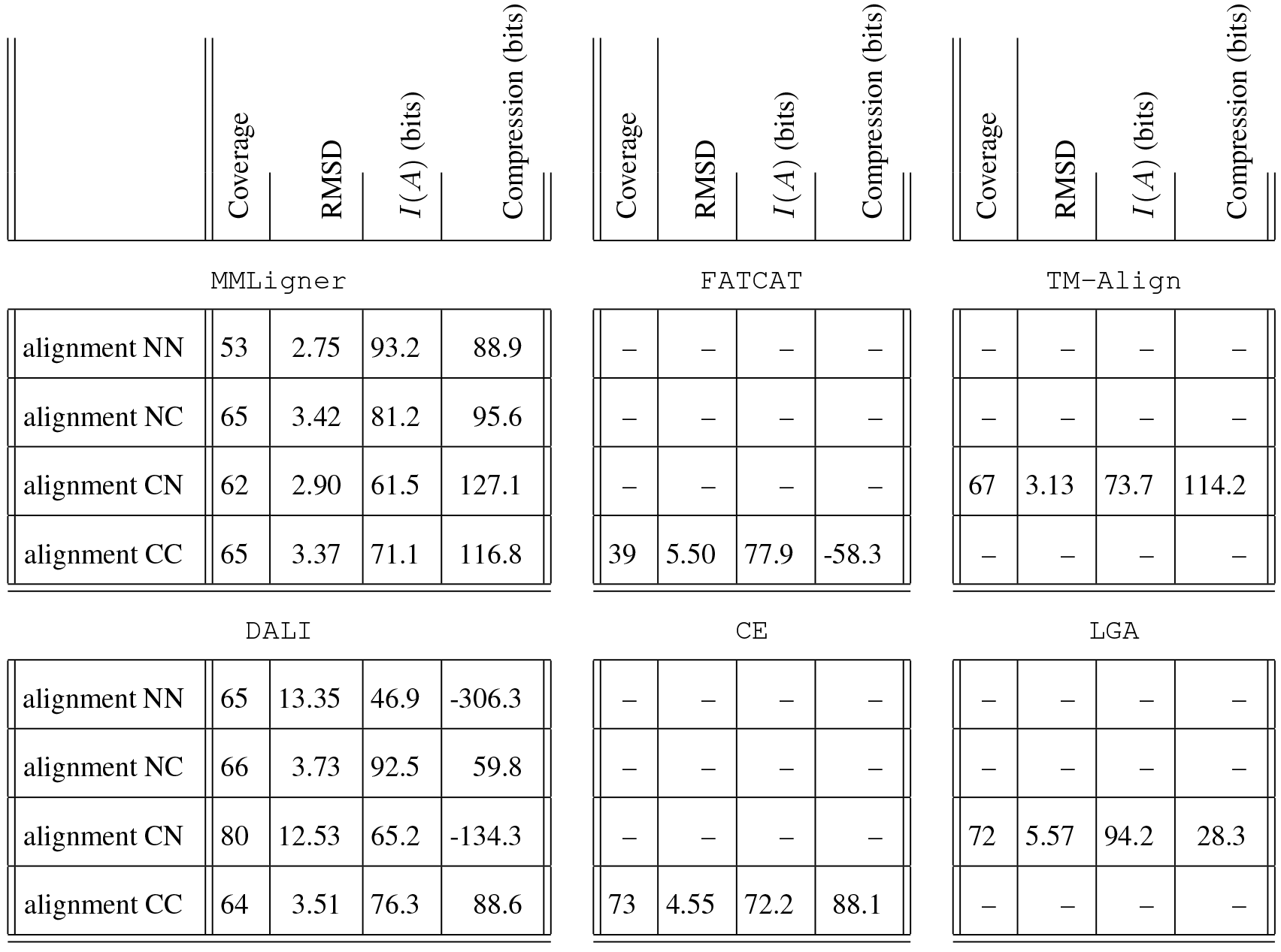
Assessment of structural alignments produced by `MMLinger`, `DALI`, `FATCAT`, `CE`, `TM-Align`, and `LGA` on the pair of calcium-binding domains, `d2sasa`_ and `d1jfja`_. Alignment NN denotes the alignment of N-terminal subunit of `d2sasa`_ (residues 1 to 99) with the corresponding N-terminal subunit of `d1jfja`_ (residues 1 to 70). Alignment CC denotes the alignment of C-terminal subunit of `d2sasa`_ (residues 100 to 184) with the corresponding C-terminal subunit of `d1jfja`_ (residues 71 to 134). (Similar definitions for the Alignments CN and CC.) The Coverage column gives the number of residue-residue correspondences reported by the respective alignment programs. The RMSD column gives the root-mean-squared-deviation after best superposition in Aunits. The *I*(*A*) column gives the measure of alignment (descriptive) complexity in bits. The compression column gives the difference between the null model message length and the `I-value` for each alignment. We use ‘-’ when no alignment is reported by any program.

**Figure 3:**
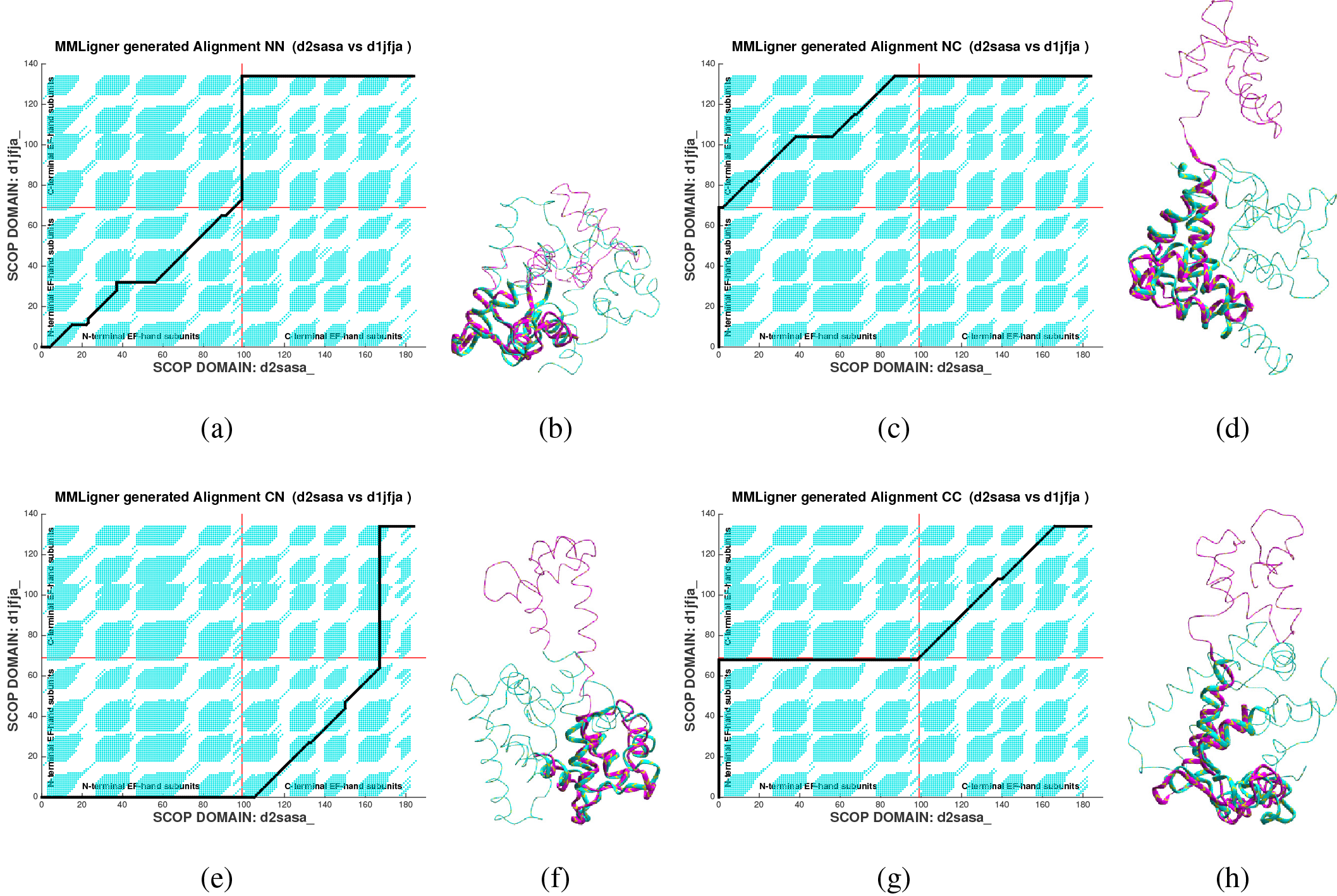
Four alignments identified by `MMLinger` when aligning the two calcium-binding protein domains `d2sasa`_ and `d1jfja`_. These alignments are denoted here as NN, NC, CN and CC. Alignment NN denotes the alignment of N-terminal subunit of `d2sasa`_ (residues 1 to 99) with the corresponding N-terminal subunit of `d1jfja`_ (residues 1 to 70). Alignment CC denotes the alignment of C-terminal subunit of `d2sasa`_ (residues 100 to 184) with the corresponding C-terminal subunit of `d1jfja`_ (residues 71 to 134). (Similar definitions for the Alignments CN and CC.) (a,c,e,g) Identified alignments (NN, NC, CN, CC respectively) shown as paths from source (bottom-left) to sink (top-right) on a 2D dot plot of MFPs found during the search in the backdrop. (b,d,f,g) Superpositions of `d1jfja`_ on d2sas_ using the identified NN, NC, CN and CC alignments respectively. (**Supplementary Movies S2(a)-S2(d))**).

**Table 2:** Performance of various structural alignment programs on the when aligning `1EUD-A` with `1CCW-A`. The Coverage column gives the number of residue-residue correspondences reported by the respective alignment programs. The RMSD column gives the root-mean-squared-deviation after best superposition in Aunits. The *I*(*A*) column gives the measure of alignment (descriptive) complexity in bits. The Compression column gives the difference between the null model message length and the `I-value` for each alignment. We use ‘-’ when no alignment is reported by any program.

| | Coverage | RMSD | $I(A)$ (bits) | Compression (bits) | | Coverage | RMSD | $I(A)$ (bits) | Compression (bits) | | Coverage | RMSD | $I(A)$ (bits) | Compression (bits) |
| --- | --- | --- | --- | --- | --- | --- | --- | --- | --- | --- | --- | --- | --- | --- |
| MMLigner |  |  |  |  |  | CE |  |  |  |  | FATCAT |  |  |  |
| d1euda1 vs. d1ccwa_ | 98 | 3.17 | 111.4 | 108.2 |  | - | - | - | - |  | - | - | - | - |
| d1euda2 vs. d1ccwa_ | 118 | 3.15 | 120.0 | 146.7 |  | 119 | 3.26 | 123.6 | 89.6 |  | 125 | 4.44 | 114.1 | 61.0 |
| d1euda1 vs. d1euda2 | 94 | 3.36 | 111.2 | 109.6 |  | 97 | 3.39 | 112.8 | 97.3 |  | 105 | 5.79 | 102.3 | -14.3 |
| DALI |  |  |  |  |  | TM-Align |  |  |  |  | LGA |  |  |  |
| d1euda1 vs. d1ccwa_ | 111 | 3.87 | 116.1 | 69.1 |  | - | - | - | - |  | - | - | - | - |
| d1euda2 vs. d1ccwa_ | 125 | 3.77 | 139.9 | 91.4 |  | 125 | 3.45 | 146.8 | 68.6 |  | 122 | 6.19 | 187.4 | -5.2 |
| d1euda1 vs. d1euda2 | 122 | 11.4 | 93.3 | -137.7 |  | 99 | 3.43 | 131.8 | 69.0 |  | 112 | 11.11 | 108.4 | -88.4 |

MMLigner and `DALI` report both alignments with the rest of the programs reporting only one of the two (Table 2). For `d1euda2 vs. d1ccwa`_, `LGA` produces the worst alignment overall, even when evaluated across all four criteria, and it is rejected according to our information measure, since there is a loss of compression of 5.2 bits, compared to the null model message length. While `DALI`, `FATCAT` and `TM-Align` produce similar and, overall, largest coverages, `TM-Align`’s RMSD is smaller. However, inspecting the alignment complexity 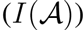, we find that `TM-Align`’s alignment complexity is significantly worse than others. As before, on manual inspection, we notice that `TM-Align` aligns many singleton or pairs of residues in regions of dissimilarity just because the chains drift together closely in space. As a result, `TM-Align` yields alignments with higher coverage (than the other programs), although with acceptable RMSD values, resulting in more complex alignment hypotheses (measured using 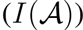. `DALI` yields an alignment with similar coverage and RMSD as `TM-Align` but with slightly less complexity. Both `MMLinger` and `CE` produce similar alignments, where differences can be attributed to the different levels of conservatism of their respective objective function.

Aligning the two domains of Succinyl-CoA synthetase, d1euda1 vs. `d1euda2` (third row of Table 2), reveals their structural self-similarity. `MMLinger`, `CE`, and `TM-Align` produce satisfactory alignments with minor differences that can be attributed to differences in the objective functions being optimised. Qualitatively, these alignments are similar. The other aligners, `DALI`, `FATCAT`, and `LGA`, produce very poor quality alignments.

In summary, `MMLinger` has the ability to identify alternative alignments for the same structural pair when they exist, and it does so consistently compared to other programs. Most programs are designed to report only one alignment. Comparison between oligomeric proteins and protein complexes will require programs to report alternative alignments. With the increasing popularity of cryo Electron microscopy in experimental protein structure determination, it is expected that the deposited structures are often large protein complexes. We therefore feel that this feature of `MMLinger` will be useful in treating the emerging datasets.

## Discussion

Several comparative studies and reviews have emphasized the inadequacy of conventional methods for protein structural alignment.^7–10^ Although many programs are guided by good biological insights, their mechanisms for assessing alignment quality are unsatisfactory and lack a proper statistical foundation. These programs define scoring schemes by combining a small number of alignment-quality measures in *ad hoc* ways. The current proliferation of structural alignment programs arise from the different ways these criteria are combined^8^.

To escape from this dilemma, we developed a new and objective approach to assessing structural alignment quality, using principles from the fields of information theory and statistical learning. The structural alignment problem is evaluated within the rigorous framework of statistical inductive inference, where alignments are hypotheses of structural relationships among observed coordinate data. Approaching the alignment problem this way has significant advantages, as the trade-off that arises when balancing the conflicting objectives of hypothesis complexity and fidelity with the observed data has to be addressed comprehensively, and without weighting any surrogate criteria. Furthermore, this treatment results in a rigorous statistical framework for discriminating between different structural alignments with comparable statistical properties, and for rejecting alignments that are not statistically significant (**Supplementary Note S1**).

Developing software to implement this information-theoretic framework has brought many challenging technical considerations, especially when designing and implementing viable statistical models of encoding to compute the various message length terms, 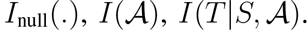. Using these, we developed a search heuristic to find meaningful structural alignments. This produced `MMLinger`, a reliable pairwise alignment program that is publicly available http://lcb.infotech.monash.edu.au/mmligner. Benchmarking its performance in comparison with other current alignment programs demonstrated that `MMLinger` is superior in consistently identifying meaningful structural alignments. The strength of the algorithm derives from its balancing hypothesis complexity 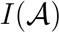 in bits) and fidelity with the observed data 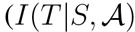 in bits). Both qualitative and quantitative inspection of the resulting alignments reveal that `MMLinger` consistently identifies statistically significant alignments, avoids pairing up random residue-residue correspondences, and finds different alignments for the same structural pair when they exist.

## Acknowledgements

Authors acknowledge Parthan Kasarapu for assistance on directional distributions that back our null model encoding.

## Supplementary notes for “*Statistical inference of protein structural alignments using information and compression*”

### S1 Bayesian framework of I-value and its statistical properties

The principles and main features of the Minimum Message Length (MML) framework are described in the main text. To summarise, given a structural alignment as a one-to-one correspondence (denoted as 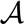) between coordinate data of a pair of protein structures (denoted by 〈*S*, *T*〉), `I-value` estimates the Shannon *information* content^1^ (denoted as 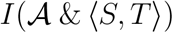 as the negativelogarithm of the joint probability of the alignment hypothesis 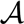 and the data 〈 *S*, *T*〉:

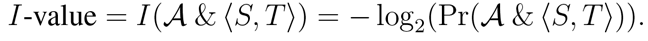

From product rule of probabilites over the events 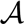, *S* and *T*, we have

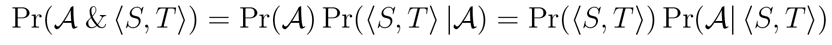

where 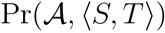 is the joint probability of alignment 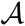 and the structural coordinates in *S* and *T*, 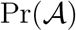 is the prior probability of the alignment 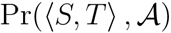 is the likelihood, Pr(〈 *S*, *T*〉) is the prior probability of the structural coordinates in *S* and *T*, and 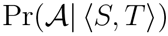 is the posterior probability of the alignment.

This product rule can be restated in Shannon’s information^1^ terms by applying negative logarithm on both sides:

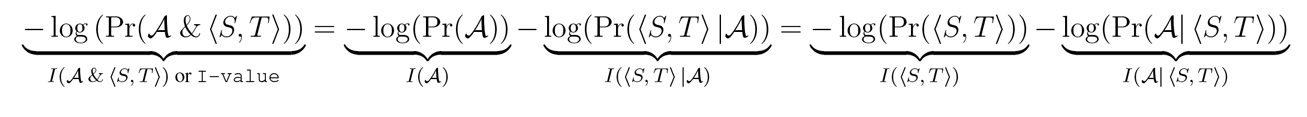

As mentioned in the main text, 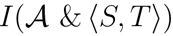 or `I-value` is computed as a length of a two-part message:

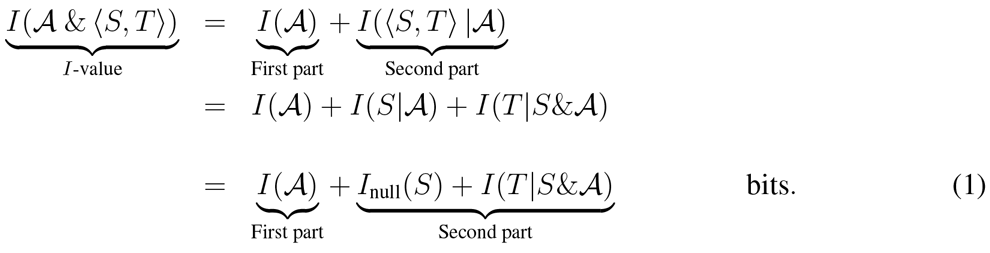

The `I-value` measure has several desirable statistical properties:

1. The `I-value` varies according to the posterior probability of 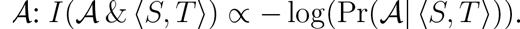
2. The difference between the `I-value`s of any two competing alignments, say 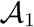 and 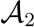, gives the log-odds posterior ratio:

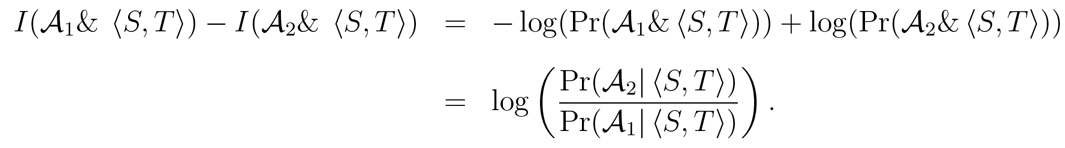 This property makes the comparison and selection of competing alignments statistically robust.
3. This framework provides a natural null hypothesis test. If the `I-value` of an alignment 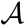 is worse (longer) than that of the null model encoding of the structural coordinates, then 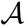 must be rejected. That is, reject 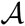 if 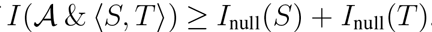. See Section S2 for details.

The computation of `I-value` involves message length terms such as 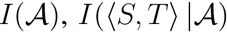, *I*_null_(*S*), *I*_null_(*T*) *et cetera* rely on probabilistic models of encoding, described below.

### S2 Computation of Null model message length: *I*_null_(*S*) and *I*_null_(*T*)

Kasarapu and Allison (2015)^2, 3^recently proposed an MML-based unsupervised learning method to infer a probabilistic mixture model using directional probability density functions. In partic-
ular, they looked at three-dimensional (3D) von Mises-Fisher and Kent probability distributions wrapped on a 3D unit sphere^2,3^. These mixture models were inferred on the empirically observed directions ^†^ of C*_α_* coordinate data from wwPDB. (See http://lcb.infotech.monash.edu.au/kent-mixture-modelling/.).

A mixture model is a probability density function of the form:

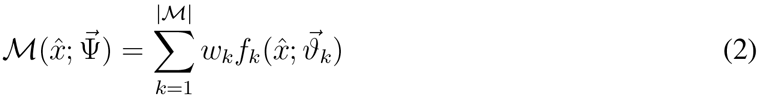

where, 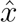 is the random variable denoting a direction, 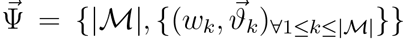 gives a vector of parameters of the mixture model containing 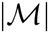 component directional probability density functions, 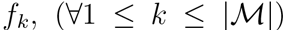, with their respective component probabilities *w_k_* (note, 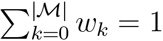 and component parameters 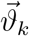.

This work employs the Kent mixture model inferred by Kasarapu (2015)^3^ on protein coordinate data. As a demonstration of the fidelity of their inferred mixture model, Fig. S1 shows the empirical distribution of C*_α_* directions (left frame) along with the distribution of randomly sampled directions from the mixture model of Kasarapu (2015)^3^ (right frame). The similarity between the two distributions clearly suggests that the mixture model faithfully characterises, into a parametric form, the empirical distribution of C*_α_* directions.

To describe the computation of the null model message length terms using the 23-component Kent mixture model^3^, *I*_null_(*S*) and *I*_null_(*T*), consider an arbitrary chain *C* containing *n* successive C*_α_* coordinates 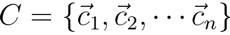. The length of the null model message we use to transmit this chain is given by:

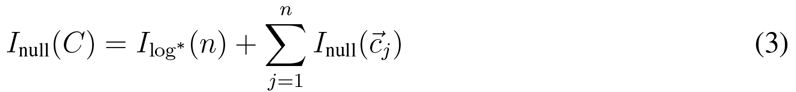

where term *I*_log*_ (*n*) represents the number of bits needed to transmit number *n* over a log* integer code.^4^ The term 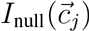 represents the number of bits needed to transmit a single 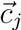 using the 23-component Kent mixture model whose computation is described below.

**Figure S1:**
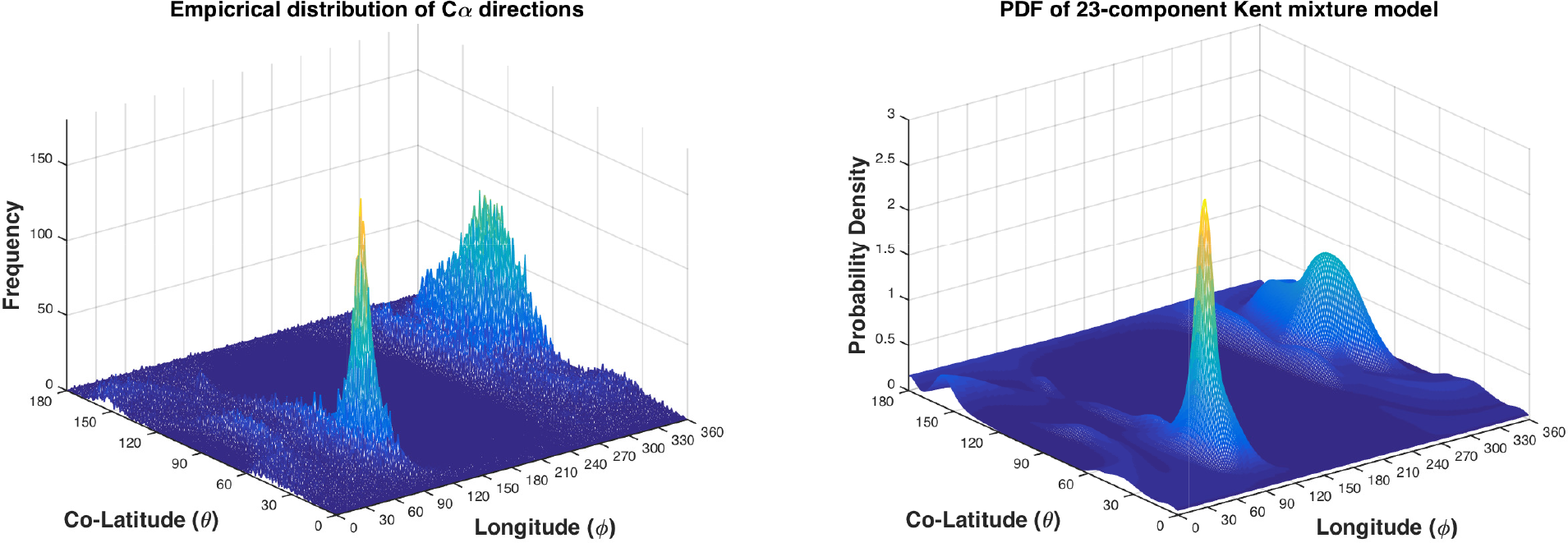
Fidelity of the 23-component Kent mixture model of Kasarapu (2015).^3^ Left frame: The empirical distribution of canonical directions of C*_α_* atoms from a set of 1,802 SCOP domains. Each observed direction is represented here using the spherical coordinates (*θ, ϕ*) denoting the spherical coordinates (also co-latitude and longitude). Right frame: The distribution of randomly sampled directions drawn from the probability distribution defined by the 23-component Kent mixture model.

Let *r_j_* and 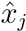 be the spherical coordinates, radius and direction (co-latitude and longitude), respectively of any coordinate 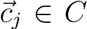. The null model message length to transmit 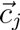 is given by 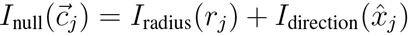 bits, that is, by the sum of the code lengths required to transmit the radius and the direction respectively. The transmission of the radius relies on the observation that the partial double bond character of the peptide bond in proteins imposes a strict constraint on the distances between successive C*_α_* atoms. Hence, each *r_j_* is the distance between C*_α_* atom 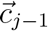 and 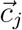, and can be encoded efficiently using a normal distribution 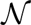 with the parameters *μ* = 3.8Å and standard deviation of *σ* = 0.2Å.^4^ This code length is computed as *I_radius_*(*r_j_*) = 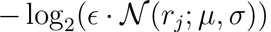.

The receipt of *r_j_* by the receiver reduces their uncertainty of the position of 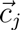 to lie anywhere on the surface of a sphere with radius *r_j_* centered at 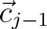. However, the direction 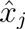 of 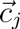 is still unknown to the receiver and must be transmitted so that the receiver can decode 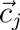 without loss of information. Each 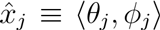 pair is encoded and transmitted in the context of the preceding three transmitted coordinates^‡^ using the 23-component Kent mixture model, denoted as 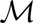 as shown in Equation 2. The code length to encode the direction is computed as 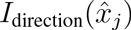 = 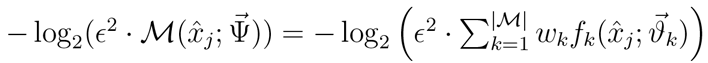.

**Time Complexity of computing the null model message length** The computation of the null model message length for any chain of C*_α_* coordinates with n atoms requires *O*(*n*) effort. This is easy to see: the computation of each 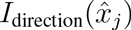 requires computing the likelihood of 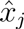 given each component density function in the null mixture. The mixture model^3^ contains a constant 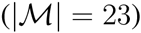 number of component Kent probability density functions. So the likelihood of each 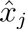 can be computed in constant time. Similarly, the computation of *I*_radius_(*r_j_*) of each C*_α_* atom is also a constant time operation. Therefore, over all *n* atoms, the computation of the null model message length of a given chain is *O*(*n*).

### S3 Computing relationship model messagelength: 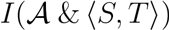

As described in the main text, the relationship model message is encoded over two-parts. In the first part the alignment 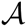 is transmitted as a string over `match (m), insert (i), delete (d) states`, followed by the coordinate data of the structures, 〈*S*, *T*〉, given this alignment relationship. The length of this two-part message is shown in Equation 1. The details of encoding required for the two parts are explained under their respective subheadings.

**Computing the first part message length**, 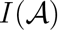: A three-state probabilistic model for alignment encoding is used here, described originally by Allison 1992^5^ to encode alignments of macromolecules, specifically over the match (*m*), insert (*i*), and delete (*d*) states. The communication of an alignment 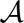 from transmitter to receiver can be achieved by first sending its length, 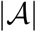, over the log* integer code, followed by sending each state symbol over a probability distribution modeled using a first-order Markov model. Such a model permits 9 possible state transitions: `mm, im, dm, mi, ii, di, md, id, dd`. Associated with each possible state transition is a transition probability: `Pr(mm), Pr(im), Pr(dm)`, and so on. These probabilities are subject to the total probability and symmetry constraints: `Pr(mm) + Pr(mi) + Pr(md) = 1, Pr(im) + Pr(ii) + Pr(id) = 1, Pr(dm)+Pr(di) + Pr(dd) = 1, Pr(mi) = Pr(im) = Pr(md) = Pr(dm), Pr(dd) = Pr(ii), and Pr(id) = Pr(di)`.

While these probabilities can be computed for any given alignment string on the transmitter’s side, the receiver however will need to know these values in advance to decode the alignment string. An adaptive code similar to the one described by Wallace and Boulton^6^ can be used to construct a decodable message over this first-order Markov model. This approach requires maintaining only 4 counters (`cntr1,…,cntr4`), one for each *distinct* probability term. These four counters are initialized to 1. The first state symbol (eg: ‘`m`’) is transmitted with a uniform probability of 1/3. Subsequently, each state symbol (eg: ‘`i`’) is transmitted over the probability that can be computed by dividing the counter pertaining to the current state transition (eg: ‘`m`’) with the sum total of all counters starting with the previous state symbol (eg: ‘`mm`’ or ‘`mi`’ or ‘`md`’). Note that both the transmitter and receiver can encode and decode (respectively) the state symbols over such a transmission. Once the current state symbol is transmitted, the counter pertaining to this state transition is incremented by one.

The code length to encode a state transition is the negative logarithm of its estimated probability. Summing up over all state transitions in the alignment string, and adding to it the code length required to transmit the size of the alignment over an integer code results in the estimation of 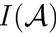 used in this work. (The symmetry constraints make the formulation of 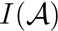 into a closed-form formula rather inconvenient. Nevertheless, the computation of 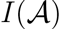 is computationally efficient as discussed below.)

**Time Complexity of computing** 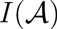 Assume |*S*| = *m* and |*T*| = *n*. Without loss of generality, assume also that *m* ≤ *n*. The maximum string length of any possible alignment 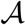 between *S* and *T* is *m* + *n* ≤ 2*n* = *O*(*n*). Using the adaptive code the computation of 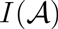 is efficient as it requires maintaining a fixed set of counters and, for each symbol in 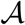, dividing two numbers and performing a negative logarithm of that ratio. Therefore, the computation of 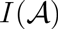 is *O*(*n*).

**Computing the second part message length**, 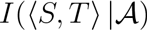: The transmission of the coordinates of *S* and *T* using the alignment information forms the second part of the message. To achieve this, the coordinates in *S* are transmitted over the null model approach described in Section S2 over a message that is *I*_null_(*S*) bits long. After decoding this information the receiver knows both 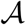 (from receiving the first part) and the coordinates of *S*. The goal now is to transmit the coordinates of *T* based on this available information.

Given 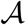 and *S*, each C*_α_* in *T* is either aligned with some C*_α_* in *S* or remains unaligned. Therefore, *T* can be partitioned into successive monotonic blocks alternating between aligned and unaligned C*_α_* coordinates. By monotonic block, we mean any run of successive C*_α_*s that are all either aligned or unaligned in *T*. For example, the alignment `iiidmmmmmmim` can be partitioned in 5 blocks (delimited by ‘`|`’ symbols) as `iii|d|mmmmmm|i|m`, where the blocks of i’s represent the unaligned coordinates of *T* while the blocks of m’s represent the coordinates in *T* that are aligned with corresponding coordinates in *S*. (Note that the blocks containing `d`’s can be ignored since they represent the unaligned coordinates in *S*, which have already been transmitted, and for this part of the transmission we are only interested in the coordinates of *T*.)

The unaligned coordinates of *T* (i.e., each chain of successive coordinates that forms any block of `i`’s in the alignment) are transmitted using the null model method described in Section S2. Thus, these unaligned coordinates offer no compression with respect to the null model transmission. Let us denote the code length to transmit these unaligned coordinates in *T* as 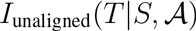.

The source of compression in this scheme potentially comes from encoding the aligned coordinates in *T*, building on the information of their corresponding coordinates in *S*. Below, we propose an efficient encoding method for such coordinates.

In this work, the estimation of the code length term, 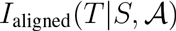, to transmit the aligned coordinates of *T* given *S* and 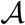, is achieved using a mixture of two encoding schemes. One takes into account the *global* spatial similarity of *positions* of aligned coordinates between *S* and *T*, while the other takes into account the *local* similarity in the *directions* of the coordinates. Appropriately, we call these two models the global and local models of encoding, respectively.

**Encoding the aligned coordinates of ***T*** using the global model**:To compute the code length of stating any 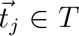 that is aligned with a 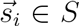 over the global model, the following procedure is employed. The two structures are first superposed based on their coordinates defined by alignment 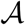. Subsequently, the set of residuals (norm of the error vectors) {*δ_ij_*}’s between each *s_i_* and *t_j_* is transmitted over a chi-squared distribution (with three degrees of freedom) using the MML approach of^7^. Further, the set of distances {*r_j_*}’s between successive *t_j-1_* and *t_j_* is transmitted using the method described in Section S2 (see Fig. S2).

**Figure S2:**
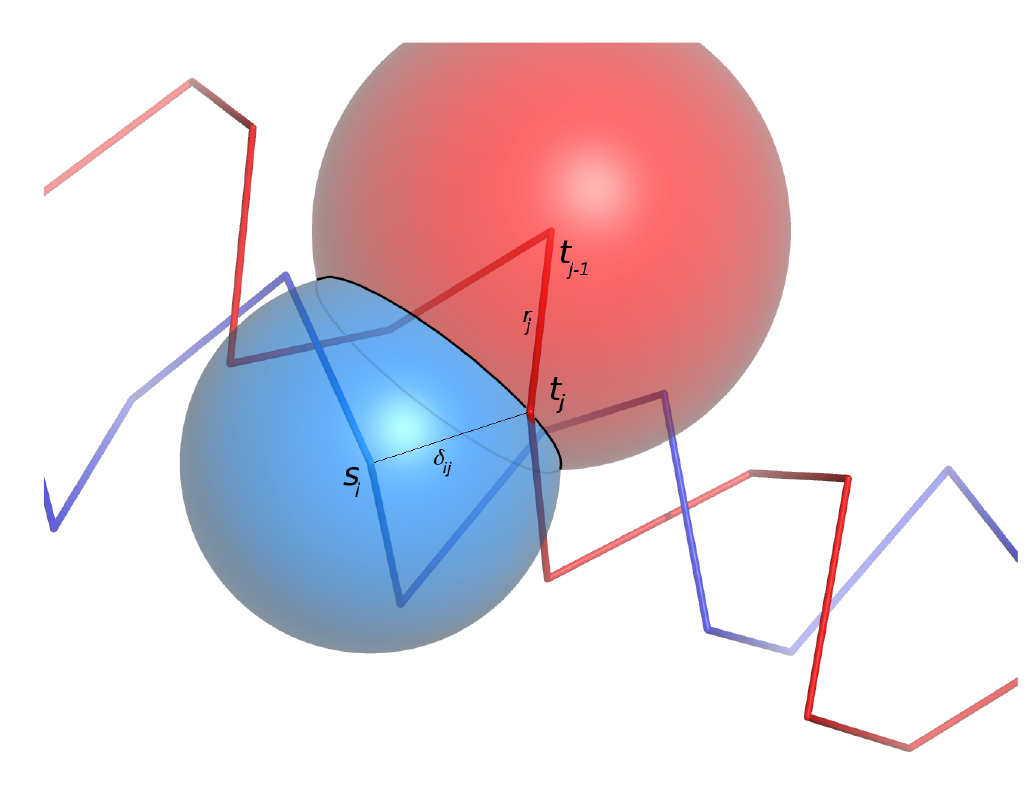
An illustration of the global alignment relationship model to transmit the coordinate of *t_j_* as a deviation with respect to *s_i_*. The receiver is sent *δ_ij_* and *r_j_* terms (see main text) which reduces the uncertainty of *t_j_* to be on a circle derived by intersecting spheres centered at *s_i_* and *t_j-1_*.

The information of *δ_ij_* and *r_j_* permits the construction of two spheres centered at 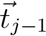 and at 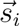. These two spheres will intersect in a circle^§^, reducing the uncertainty of *t_j_* to lie on the circle of intersection. Given this, the coordinates of 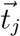 can be transmitted over a uniform distribution on the circle (see Fig. S2).

**Encoding the aligned coordinates of *T* using the local model**: The first three coordinates in every aligned block of *T* are encoded using the global model described above. For the remaining of matches in each mathced block, the following local encoding method is employed.

Recall from Equation 2 in Section S2, that the mixture model is of the form 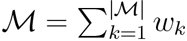. 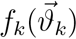, where *w_k_* is the associated probability of the *k*th component (which in turn is a Kent directional probability distribution.) If no (extra) information were available about 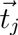, then the transmitter would have no choice but to encode it using the null model. However, extra information is available in the form of the correspondence of 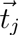 with 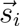. Thus, the probabilities of each component can be updated based on this relationship. Specifically, the null component probabilities (*w_k_*’s) can be systematically updated given the expectation provided by the alignment that 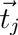 is near 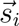.

Bayes theorem allows updating all *w_k_*’s given the knowledge of 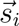. Consider the computation of the posterior probability of the *k*th component given the direction 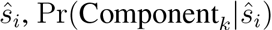:

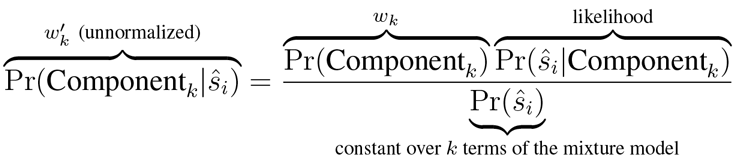

Note that Pr(Component_*k*_) is equal to *w_k_* of the null model, and 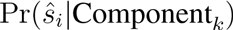 is the likelihood of the direction 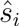 given the component probability density function, 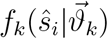. While these two can be computed trivially using the null mixture model 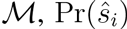 is unknown. However, each 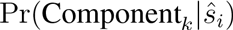 needs to be normalised (so that their sum over all components adds to 1) by dividing the numerator by 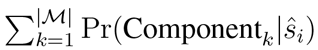, the term disappears. Call these normalised posterior weights *w′_k_*. The old *w_k_*’s are now replaced with the new *w′_k_*’s in the null mixture model 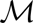 and *t_j_* can now be encoded using the updated mixture model.

**Time Complexity of computing** 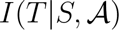 Assuming again that |*S*| = *m* and |*T*| = *n* and *m* ≤ *n*, the global model of encoding requires the superposition of the aligned coordinates between *S* and *T*. The maximum number of aligned coordinates in any 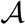 is bounded by *m*. The computational effort to find the best superposition of two corresponding vector sets containing *m* vectors is *O*(*m*).

For the global model the estimation of code length for *δ_ij_* and *r_j_*, as well as the encoding over the circle of intersection (between spheres) takes constant time. Over all aligned coordinates, the time complexity to estimate the message length over the global model takes worst-case *O(m)* time as the number of matches is bounded by *m*. This is the same for the local encoding as each aligned 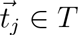 requires the computation of the re-weighted probabilities (*w′_k_*’s) that take a constant 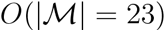) effort.

On the other hand, the encoding of unaligned coordinates of *T* (encoded using the null model) is *O(|T|)* = *O*(*n*) in the worst-case. Therefore, the estimation of `I-value` given the coordinates of *S* and *T* and any alignment between them, 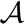, takes *O*(*n*) time overall.

### S4 MMLigner search heuristic

`MMLigner`’s search method is carried out in two phases. In the first phase, seed structural alignments are quickly generated using a deterministic approach that efficiently clusters and consistently assembles well-superposable fragment pairs for the two given structures *S* and *T*. These seed alignments are refined in the second phase using the `I-value` criterion over a heuristic search.

**Phase 1: Generating alignment seeds** *Identification of a library of maximal fragment pairs (MFPs)*. Given a pair of structures *S* and *T*, we first identify all well-superposable, maximally-extended fragment pairs (MFPs) that fit within a threshold of RMSD (= 2.5 Å). This results in a library of MFPs for the given pair of structures being aligned.

*Filtering the library of MFPs*. The library of MFPs is then filtered to contain only those MFPs that can be jointly superposed with at least two other MFPs in the set. Specifically, every pair of non-overlapping MFPs are jointly superposed under the threshold of RMSD of (= 3 Å). Any MFP that does not superpose within this threshold will be discarded since it shows no evidence of being spatially consistent with other MFPs in the set. This eliminates MFPs that are locally but not globally meaningful.

For each pair that jointly superposes within the threshold of RMSD, we look to extend the superposition using yet another (non-overlapping) MFP. Any pair of MFPs that is not extendible and whose combined length is < 15 residues is again discarded. This further prunes the original library of MFPs.

We note that, if carried out naively, this filtering step can be computationally expensive. However, we are able to achieve an exhaustive and extremely efficient joint superpositions over all pairs of MFPs (and their further extensions to triples) by benefiting from our recent work that has identified the sufficient statistics for orthogonal least-squares superposition of vector sets ^8^. To do this, when the original library of MFPs is computed, we store the sufficient statistics of the superposition in each MFP. The RMSD of joint superposition of pairs of MFPs as well as their sufficient statistics can be computed as a constant time update using the sufficient statistics of individual MFPs.^8^ Similarly, extensions of pairs of MFPs to triples can also be updated in constant time. As a result, the identification of MFPs and the pruning can be carried out exhaustively in seconds.

*Clustering the filtered set of MFPs*. The aim of this step is to partition the filtered set of MFPs into groups (or clusters) of related MFPs so that a seed alignment can be explored within each cluster.

Our iterative clustering heuristic proceeds as follows. First, the filtered set of MFPs is sorted in decreasing order of length (in terms of number of residue pairs in each MFP) and the longest MFP in the filtered set is assigned as the initial singleton cluster. The iterative process of clustering then starts by traversing the remaining sorted list of filtered MFPs, starting from the longest. For each MFP in the list, we attempt to add it to any of the existing clusters. Otherwise, a new cluster is created with this MFP as its singleton member. In general, the MFP can be added to a cluster if that MFP jointly superposes with at least 40% of the cluster’s existing MFPs.

At the end of this procedure, the clusters whose combined length is less than 18 residues are deleted and the remaining ones are used in the next step to identify seed alignments.

*Finding a seed alignment using the clustered MFPs*. For the MFPs represented in each cluster, a scoring matrix is computed. Each cell in this matrix represents a score for aligning a specific residue-pair between the two structures being aligned. For each triplet of MFPs in the set that superpose within the threshold of RMSD (3 Å), scores (computed from the combined lengths of MFPs involved and their RMSD) are populated in the weight matrix for the residue-pairs involved in those MFPs. A rough seed alignment using a dynamic programming algorithm described in the pairwise step of `MUSTANG`^9^.

**Phase 2: Iterative refinement of the seed alignment using `I-value` criterion** Using each seed alignment identified in the previous phase as the starting point, a series of perturbations to these alignments are carried out to identify the final alignment(s) that `MMLinger` reports. The fitness of each perturbed alignment is evaluated using the `I-value` measure based on the amount of compression achieved compared to the null model.

This approach that is similar to the Expectation-Maximization (EM) technique common in statistical learning. Each alignment is efficiently represented as a vector of blocks (where each match block stores the start indices in *S* and *T* of that match block, followed by the length of the block). Until the method converges (or reaches a maximum of iterations), the current *best* alignment at each iteration is operated upon by performing the following primitive permutation operations, on each of the blocks.

**ExtendMatchBlock(***i*, *l*, **direction)**: This search primitive tries to extend the *i*-th block in an alignment by *l* residues either on the left side or on the right side (depending on the given direction) of the match block. That is, it tries to create new matches out of the deletes and inserts (if any) flanking the specified block in the specified `direction`. The number of new matches is limited by min(‖*inserts*‖, ‖*deletes*‖).

**ShrinkMatchBlock(***i*, *l*, **direction)** This primitive tries to shrink the *i*-th match block by *l* residues either on the left or right of the match block. That is, it tries to remove macthes by creating new insertions and deletions (*indels*) flanking the current block in the specified direction. The number of new indels is limited by ‖*matchblock*‖.

**ShiftMatches(***i*, *l*, **direction)**: This primitive tries to shift *l* aligned residues from the start of the current block to the end of the previous one, or vice versa. This can only be done when the gap separating two successive match blocks is monotonous (that is, the gap is either entirely inserts, or entirely deletes). Upon this perturbation, the gap length between the successive blocks remains unchanged, but the size the specified match block either increases or decreases by *l*. The shift size, in turn, is limited by the size of the previous or current block, depending on the direction of shift.

**SlideMatchBlock(***i*,*l*, **direction)**: This primitive tries to change the residue-residue correspondences of a block by moving (or *sliding*) *l* residues in *S* left or right relative to *T*. Note that the number of correspondences in a block remains constant. The size of the shift is limited by the size of the block being operated upon plus the number of gaps available in the direction of the shift.

**RealignClosest(***i*): This primitive perturbs the current alignment by realigning the (subsets of) residues in *S* and *T* around a specified match block. Before this perturbation is explored, *T* is superposed on *S* based on the current (full) state of the alignment. For any *i*-th match block, consider the subsets of residues in *S* and (transformed) *T* covering the specified match block, and its left and right flanking gap region. A Euclidean distance matrix is computed for each residue-residue pairs within this subsets. The residue-pairs within these subsets that exceed the maximum match distance observed with the current match block are ignored from the realignment. For the permissible residue pairs, a Needleman-Wunsch^10^ dynamic programming algorithm is used to realign the residues around this specified region.

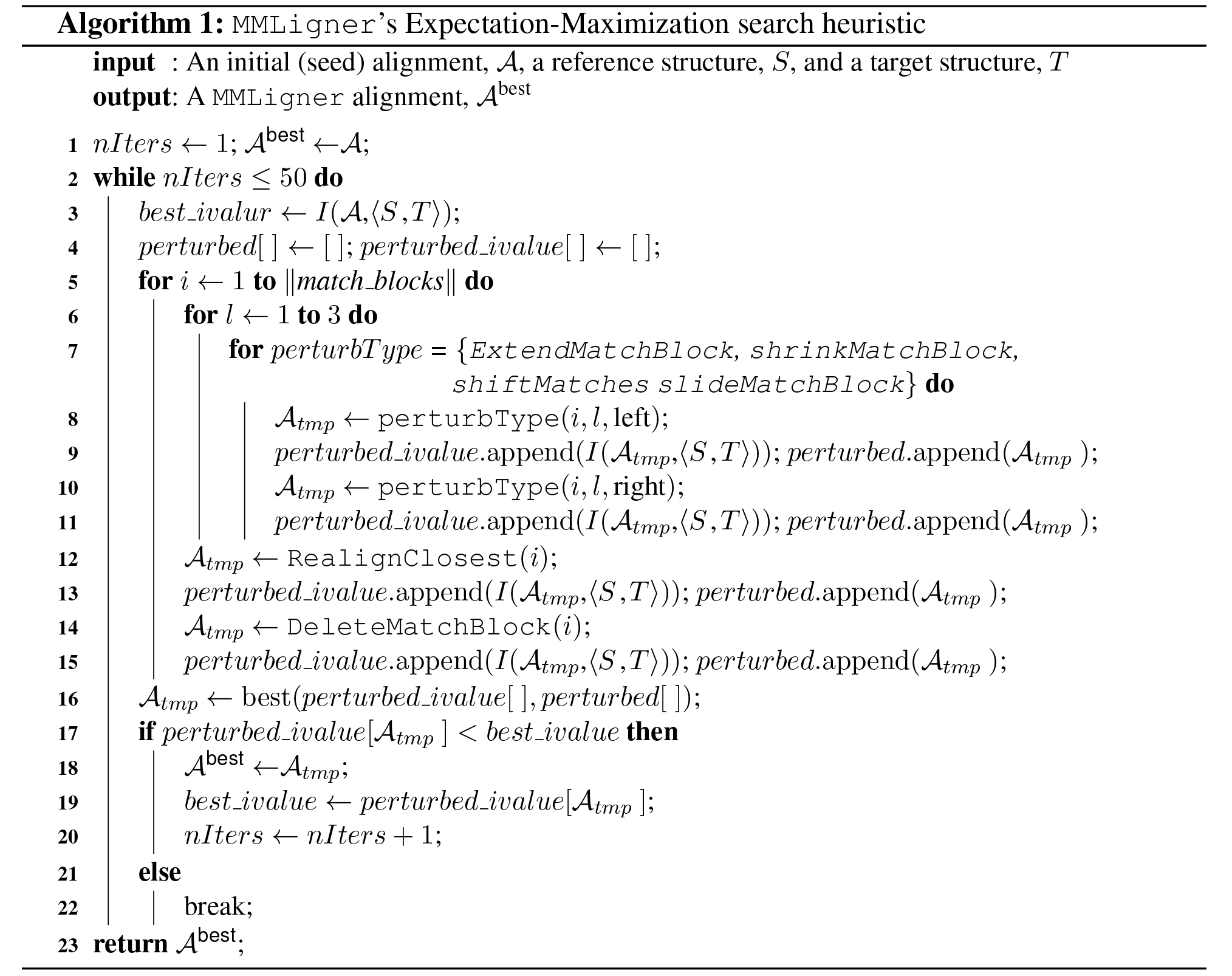

**DeleteMatchBlock(***i*): This primitive perturbs the current alignment by deleting the entire match block and replacing it with appropriate inserts and deletes.

**Search using the above primitives**: Any given seed alignment is refined using the above primitives over an Expectation-Maximization algorithm defined in Algorithm ??. At each iteration, the algorithm attempts to exhaustively perturb each match block using the above perturbation primitives and greedily chooses the best perturbation using the `I-value` criterion. This continues until either the alignment converges (it always will but might take long) or reaches the maximum number of iterations. This behaviour is intended to ensure the algorithm explores the local space thoroughly around the (reasonably good) starting point provided by the seed alignment of `MMLinger`.

### S5 Selecting the SCOP domain dataset used for software benchmarking

In the results section of the main text, we benchmark various alignment program on a dataset containing 2500 structural domain pairs selected randomly from SCOP domain database.^11, 12^No two domains in our dataset share more than 40% sequence identity.

The dataset is identified using the following procedure. ASTRAL SCOP 40^12^ domains are used and separated out into buckets depending on their sizes (number of residues). Two domains in the same bucket differ no more than 50 residues in their lengths. The SCOP hierarchy for all the domains within each bucket is recorded. A pivot domain is randomly chosen from the entire ASTRAL SCOP 40 collection. Assume that this pivot domain falls with the i-th bucket. Using the SCOP hierarchy in this bucket, we select:

- one domain (randomly) that belongs to the same SCOP Family as the pivot
- one domain (randomly) that belongs to the same SCOP Superfamily as the pivot, but not the Family
- one domain (randomly) that belongs to the same SCOP Fold the pivot, but not the Family or Superfamily.
- one domain (randomly) that belongs to the same SCOP Class the pivot, but not the Family, Superfamily or Fold.
- one domain (randomly) that belongs to a different class.

This selection process is repeated until we have 500 distinct pivots and their respective five domains. This results in 2500 distinct structural domain pairs. (Dataset is available from http://lcb.infotech.monash.edu.au/mmligner).

### S6 Supplementary figure supporting Fig. 2. in the main text

`MMLigner` can be run with an option to filter out alignments for structural pairs composed of matches involving standard supersecondary structures, which nevertheless yield positive compression over their respective. These are reported by `MMLinger` in its default run (see Fig. 2. in the main text). Fig. S3 shows the runs with the filter option turned on, over the same dataset used in the main text.

**Figure S3:**
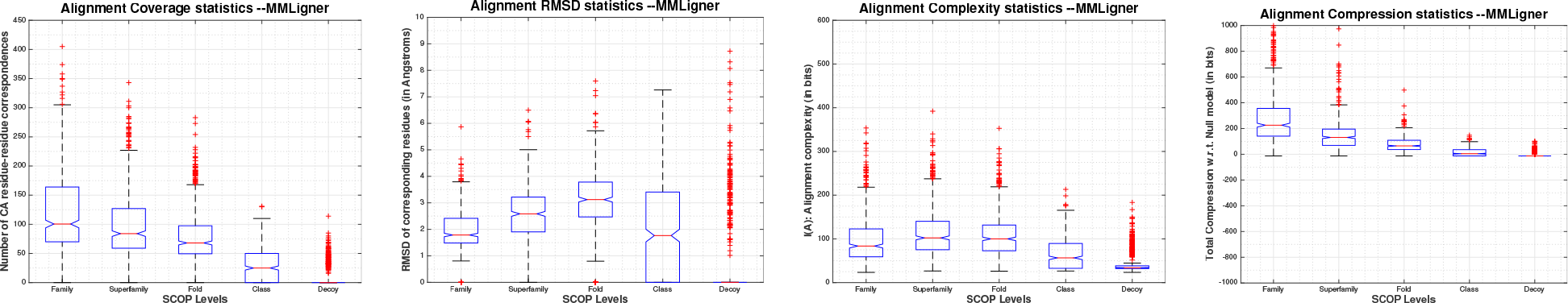
Supplementary figure to Fig. 2. in the main text. Notched Box-Whisker plots displaying the distribution of values from various criteria across 500 × 5 = 2500 alignments generated by `MMLinger` with a filter option turned on (default is off; see results in Fig. 2. in the main text). The filter option ignores any alignment(s) returned by `MMLinger` that, although yielding positive compression, is composed of matches involving common supersecondary structures.

## Acknowledgements

Authors acknowledge Parthan Kasarapu for assistance on using his mixture models.

## Website

MMLigner software and all material related to this work is found at http://lcb.infotech.monash.edu.au/mmligner.

## Footnotes

† Directions are given by the spherical coordinates of C*_α_* atoms, denoting the zenith angle (or co-latitude) and azimuth angle (or longitude), measured in a canonical frame of reference.

‡ Since this encoding requires a context of three preceding residues, as a boundary case, the first three C*_α_*s are transmitted using the null model

§ Ignoring the pathological case when *t_j-1_* = *s_i_* or in the best case when *s_i_* = *t_j_*

